# A novel *hemA* mutation is responsible for small colony variant phenotype in *Escherichia coli*

**DOI:** 10.1101/2020.05.01.073478

**Authors:** Alasdair T. M. Hubbard, Adam P. Roberts

## Abstract

We identified a small colony variant (SCV) of a clinical isolate of *Escherichia coli* from Malawi following sequential *in vitro* selection in sub-inhibitory concentrations of amoxicillin-clavulanic acid and gentamicin. The SCV was auxotrophic for hemin and had impaired biofilm formation compared to the ancestral isolates. A single novel nucleotide polymorphism (SNP) in *hemA*, which encodes a glutamyl-tRNA reductase responsible for the initial step of porphyrin biosynthesis leading to the production of haem, was responsible for the SCV phenotype. We showed this phenotype was stable over multiple generations and the SNP in *hemA* resulted in a significant fitness cost to the isolate which persisted even in the presence of hemin. As *hemA* is not found in mammalian cells, and disruption of the gene results in impaired biofilm formation and a significant fitness cost, it represents a potential target for novel drug development specifically for the treatment of catheter-associated urinary tract infections caused by biofilm-producing *E. coli*.

## Introduction

Small colony variants (SCVs) are slower growing mutants observed in the study of multiple bacterial species. They have altered, smaller than normal, colony morphology which is usually due to mutations within genes involved in metabolic processes, for example in genes involved in the electron transport chain of *Staphylococcus aureus* [1]. SCVs have been shown to be responsible for multiple infections including tissue, bone and device-associated infections [2] and can result in increased resistance to aminoglycosides [3, 4] which can also be used to select for SCVs *in vitro* [5]. The mechanism of decreased susceptibility to aminoglycosides has not been well investigated but in *S. aureus* it has been shown to be due to alterations in the membrane potential and uptake [6]. Investigations in SCVs of *Escherichia coli* are relatively rare but have also been associated with mutations within proteins of the electron transport chain, including *yigP* [4], *hemB* [7] and *lipA* [8]. SCVs of *Escherichia coli* have been found to be etiological agents of urinary tract infections [9], bacteraemia [10] and chronic hip infections [7] and are therefore increasingly considered clinically relevant [11].

An important enzyme involved in the electron transport chain is HemA. HemA is a glutamyl-tRNA reductase which catalyses the initial step of porphyrin biosynthesis leading to the production of haem, which is required for respiration. Single nucleotide polymorphisms (SNP) identified within *hemA* have been found to result in a SCV in several bacterial genus, including *Salmonella* [3, 12] and *Listeria* [13,14], as well as in *E. coli* [15]. The morphology of three SCVs of *Listeria monocytogenes*, all with a common SNP present in *hemA* resulting in early termination of the protein, were rescued when grown in the presence of exogenous hemin, showing they were auxotrophic for haem [14].

Here we report the identification of a novel SNP in *hemA* resulting in a SCV phenotype of a clinical isolate of *E. coli* from Malawi which was sequentially selected for in subinhibitory concentrations of amoxicillin/clavulanic acid and gentamicin. We assessed the effect of this SNP on the fitness of the SCV compared to the ancestral isolates and, as biofilm-producing *E. coli* are common etiological agents of catheter-associated UTIs [16], we sought to determine the biofilm forming ability of the SCV.

## Materials and methods

### Bacterial strains, media and antibiotics

*E. coli* 10129 is a susceptible clinical isolate from Malawi isolated from cerebral spinal fluid [17] and *E. coli* 10129 LB_2 is an *in vitro* derived isolate with resistance to amoxicillin/clavulanic acid (AMC) [18]. Both isolates were previously whole genome sequenced and are available in GenBank under the following accession numbers: *E. coli* 10129; SIJF00000000 and *E. coli* 10129 LB_2; SAMN10963734.

All bacterial cultures were initially grown on LB (Lennox) agar (Sigma, UK) or CHROMagar™ Orientation (CHROMagar, France) for 18 hours at 37°C then subcultured into cation-adjusted Mueller Hinton Broth (CA-MHB), LB (Lennox) broth (LB) (both Sigma, UK) or iso-sensitest broth (ISO, Oxoid, UK) and incubated at 37°C, 200 rpm for 18 hours unless otherwise stated.

Gentamicin (gentamicin) were solubilised in molecular grade water (all Sigma, UK) and filter sterilised through a 0.22μM polyethersulfone filter unit (Millipore, US).

### Selection of gentamicin resistant colonies

An initial culture of *E. coli* 10129 LB_2 was diluted 1/1000 in LB plus 1 μg/ml gentamicin and incubated for 24 hours at 37°C, 200 rpm. Following incubation, the culture was serially diluted 1 in 10 to the 10^-2^ dilution and 50 μl plated out on LB agar plus 16 μg/ml gentamicin and incubated at 37°C for 18 hours. A SCV, designated *E. coli* 10129 LB_2.6 and a normal sized colony, designated *E. coli* 10129 LB_2.5 were pure cultured and stored at −80°C.

### Antimicrobial susceptibility testing

Minimum inhibitory concentration of gentamicin towards *E. coli* 10129, *E. coli* 10129 LB_2 and *E. coli* 10129 2.6 were determined using the microdilution broth method following the CLSI guidelines using CA-MHB and performed in triplicate.

### Characterisation of colony and cell morphology

*E. coli* 10129, *E. coli* 10129 LB_2, *E. coli* 10129 LB_2.5 and *E. coli* 10129 LB_2.6 were all grown on LB agar, both with and without supplementation of 20 μg/ml hemin (Sigma, UK) and CHROMagar™ Orientation agar for 18 hours at 37°C, then photographed under a Celestron MicroDirect 1080p HD Handheld Digital microscope (Celestron, US). Gram stain of each isolate was performed using a standard protocol (Sigma, UK) and visualised under a Motic Trinocular light microscope mounted with a Moticam X Lite camera (Motic, Hong Kong)

### Whole genome sequencing and bioinformatics analysis

Short-read (2 x 250 bp paired end) genome sequencing of *E. coli* 10129 LB_2.6 using the Illumina MiSeq platform, and subsequent read trimming, was provided by MicrobesNG (http://www.microbesng.uk). The genome was *de novo* assembled using SPAdes (v3.12.0, [19]) and annotated using Prokka (v1.12, [20]). The sequencing reads of *E. coli* 10129 LB_2.6 were aligned to the annotated assembled genomes of *E. coli* 10129 and *E. coli* 10129 LB_2 to identify statistically relevant SNPs, indels, duplications, and amplifications using Breseq (v0.32.0, [21]). Protein structure predictions were carried out using I-TASSER (https://zhanglab.ccmb.med.umich.edu/l-TASSER/) [22] and comparisons carried out using Tm-align (https://zhanglab.ccmb.med.umich.edu/TM-align/) [23]. Protein models used in structural comparisons were selected based on the highest C-score and TM-score.

### DNA extraction

DNA extraction was performed using the PureGene^®^ Yeast/Bact kit B (Qiagen, Germany) following the manufacturer’s instructions for extraction of DNA from Gram-negative bacteria and eluted in molecular grade water.

### PCR

All PCR was performed using 0.02 U/μl Q5^®^ High-Fidelity DNA Polymerase (New England Biolabs, US), 1x Q5^®^ Reaction Buffer, 0.5 μM primers and 200 μM dNTPs in a total volume of 25 μl and, unless otherwise stated, all PCR products were purified using the Monarch^®^ PCR and DNA Clean-up Kit (New England Biolabs, USA) following the manufacturer’s instructions and eluted in molecular grade water.

Amplification of 819 bp of *hemA* was performed the primers HemA_F (TGTCGACGTGTAACCGCACA) and HemA_R, (CCACAGCAGCAGCTTTCCGTTG) and the following protocol: Denaturation at 98°C for 30 seconds, followed by 35 cycles of denaturation at 98°C for 10 seconds, annealing at 70°C for 30 seconds and elongation at 72°C for 50 seconds, followed by a single, final extension of 2 minutes at 72°C. The entire 1257 bp of *hemA* from *E. coli* 10129 using the primers HA1_F (ATGACCCTTTTAGCACTCGG) and HA1_R (CTACTCCAGCCCGAGGCT) in a total volume of 50 μl and the following protocol; Denaturation at 98°C for 30 seconds, followed by 35 cycles of denaturation at 98°C for 10 seconds, annealing at 66°C for 30 seconds and elongation at 72°C for 13 seconds, followed by a single, final extension step 72°C for 2 minutes. A band at approximately 1 kb was cut out and DNA extracted using the Monarch DNA Gel Extraction Kit (New England Biolabs, US) following the manufacturer’s instructions. The restriction sites for the restriction enzymes KpnI were added to the 5’ ends of the 1257 bp double stranded amplicon by a second round of PCR, using the primers KpnIHA_F1 (AAAAA<u>GGTACC</u>ATGACCCTTTTAGCACTCGG) and KpnIHA_R1 (AAAAA<u>GGTACC</u>CTACTCCAGCCCGAGGCT) (Kpn1 site is underlined). Amplification was performed using the same protocol as the initial amplification of *hemA*.

### Stability of the small colony phenotype

*E. coli* 10129, *E. coli* 10129 LB_2 and *E. coli* 10129 LB_2.6 were passaged sequentially for 40 days on LB agar and incubated at 37°C. Each day, the colonies of each of the three isolates where photographed under a Celestron MicroDirect 1080p HD Handheld Digital microscope.

### Construction of complement plasmid

The resulting PCR product of the PCR amplifying the wildtype *hemA* with added KpnI sites and the recipient plasmid pHSG396 were digested with the restriction enzymes Kpn1 (New England Biolabs, US). PCR clean-up was then performed using the Monarch PCR and DNA Clean-up Kit following the manufacturer’s instructions. The digested *hemA* PCR product and pHSG396 were ligated together using T4 ligase (New England Biolabs, US), at a 3:1 insert to vector mass ratio, incubated at room temperature for 10 minutes followed by 65°C for 10 minutes.

The ligated, recombinant plasmid (2 μl) was transformed into NEB^®^ 5-alpha competent *E. coli* (New England Biolabs, US) using the following protocol: the reaction was incubated on ice for 30 minutes, then heat shocked at 42°C for 30 seconds. Following a second incubation on ice for 5 minutes, 950 μL of SOC outgrowth medium was added and incubated at 37°C, 250 rpm for 1 hour. The transformed cells were plated out onto LB agar supplemented with 35 μg/ml chloramphenicol and Isopropyl β-D-1-thiogalactopyranoside/X-gal (Fisher Scientific, USA) and incubated at 37°C for 18 hours and the plasmid was extracted using the Monarch Plasmid Miniprep Kit (New England Biolabs, US) following the manufacturer’s instructions. *hemA* in pHSG396 was Sanger sequenced by GeneWiz (Takely, UK) using commercially available M13 primers. *E. coli* 10129 LB_2.6 was made competent following the protocol set out in Chung *et al.* 1989 [24] and pHSG396 and pHSG396:hemA was transformed following the above protocol.

### Biofilm assay

Biofilm production by *E. coli* 10129, *E. coli* 10129 LB_2, and *E. coli* 10129 LB_2.6, was quantified as described previously [18]. Briefly, each isolate was initially grown in 10 ml LB, then diluted 1/1000 in M9 (50% (v/v) M9 minimal salts (2x) (Gibco, ThermoFisher Scientific, USA), 0.4% D-glucose, 4mM magnesium sulphate (both Sigma, UK) and 0.05 mM calcium chloride (Millipore, USA)) with and without supplementation with 20 μg/ml hemin. Four technical replicates of each diluted culture were added to a 96 well plate and incubated statically for 18 hours at 37°C. Following incubation, each well was washed with PBS, stained with 0.1% crystal violet and washed again with PBS. The stain was liberated from the 96 well plated with 30% acetic acid (Fisher Scientific, USA) and measured at an OD_550_. Three biological replicates of this assay were performed.

### Competitive and comparative fitness assays

Competitive fitness of *E. coli* 10129 LB_2.6 was assessed compared to *E. coli* 10129 and *E. coli* 10129 LB_2 in LB and M9 as described previously [18]. Briefly, each culture was diluted to an OD_600_ of 0.1 and then dilute 1/1000 and combined 1:1 in the appropriate media and 150 μl added to a 96 well plate and incubated at 37°C, 200 rpm for 18 hours. Bacterial density of the initial combined culture and after 24-hour incubation was enumerated by diluting the culture in PBS and 50 μl of each dilution plated out on LB agar and LB ager plus 20 μg/ml gentamicin and incubated at 37°C for 18 hours.

Fitness of *E. coli* 10129 LB_2.6, *E. coli* 10129 LB_2 with pHSG396 and *E. coli* 10129 LB_2.6 with pHSG396:hemA were compared to E. coli 10129 LB_2 comparatively. Each culture was diluted to an OD_600_ of 0.1 then further diluted 1/1000, both in LB, and 150 μl of this diluted culture was added to a 96 well plate in duplicate. As a negative control, 150 μl LB or LB with 35 μg/ml chloramphenicol and 1 mM IPTG was also added to the plate in duplicate. All samples were incubated at 37°C for 24 hours, orbital shaking at 200 rpm, in a Clariostar Plus microplate reader (BMG Labtech, Germany) measuring the OD_600_every 10 minutes. Relative fitness of each isolate was compared to *E. coli* 10129 LB_2 using BAT v2.1 [25] with absorbance values between 0.02 and 0.2 and a minimum R value of 0.9902. Both fitness assays were performed in triplicate.

### Statistical analysis

All statistical analysis was performed using the ordinary one-way ANOVA plus uncorrected Fisher’s LSD test in GraphPad Prism (v8.4.0).

## Results

### Isolation of a small colony variant

After selection of gentamicin resistant *E. coli* 10129 LB_2 derivatives following growth in the presence of sub-inhibitory concentrations of gentamicin, we noticed a small pinprick colony on the agar plate which was immediately stored at −80°C and designated *E. coli* 10129 LB_2.6. Subsequent regrowth of *E. coli* 10129 LB_2.6 confirmed the SCV morphology persisted when compared to the two ancestral strains and another normal sized isolate, *E. coli* 10129 LB_2.5, from the same selective plate (Fig. 1A). The SCV was confirmed to be Gram negative, however the cell size and shape was not observed to be different than that of the two ancestral isolates or the *E. coli* 10129 LB_2.5 strain (Fig. 1C). To ensure that the SCV was not a Gram-negative contaminant, we confirmed it was *E. coli* through growth on CHROMagar™ Orientation agar (Fig. 1B). The MIC for gentamicin for *E. coli* 10129 LB_2.6 is 8 μg/ml compared to 1-2 μg/ml for *E. coli* 10129 and 2-4 μg/ml *E. coli* 10129 LB_2.

**Figure 1:**
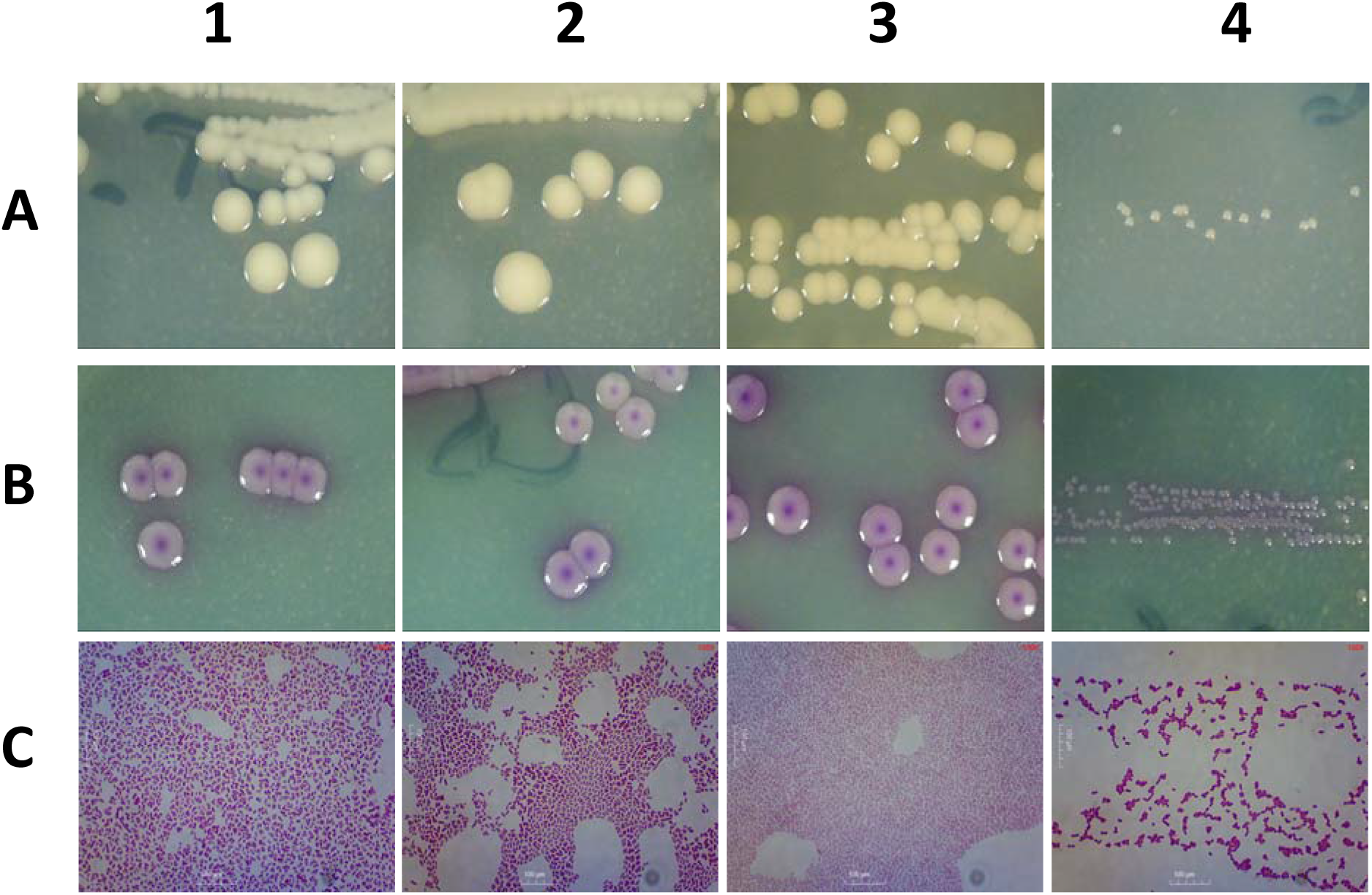
Photographs of the colonies of 1) *E. coli* 10129, **2)** *E. coli* 10129 LB_2, **3)** *E. coli* 10129 LB_2.5 and **4)** *E. coli* 10129 LB_2.6 (SCV) grown on **A)** LB agar, **B)** CHROMagar™ Orientation agar and **C)** Photographs of the Gram stain of each of the isolates.

### Mutations identified within the genome of E. coli 10129 LB_2.6

By comparing the whole genome sequence of the *E. coli* 10129 LB_2.6 to the ancestral isolates *E. coli* 10129 and *E. coli* 10129 LB_2, we identified a single novel SNP present in *hemA* in *E. coli* 10129 LB_2.6 (T>G at nucleotide 426 out of 1257). The SNP was predicted to result in a non-synonymous mutation leading to a change of phenylalanine to leucine at amino acid position 142. This SNP was confirmed to be present in *E. coli* 10129 LB_2.6 but not in the two ancestral strains through PCR amplification and Sanger sequencing of *hemA* in all three isolates (results not shown).

### Structural changes are predicted in the HemA from the SCV compared to the wildtype

The phenylalanine to leucine change at amino acid 142 is predicted to be within an α-helix that links two globular domains within the tertiary structure of the proteins, the relative positions of which are predicted to change (Figure 2). This is predicted to alter the position of the substrate binding sites for both NAD and glutamyl-tRNA, as determined by Tm-align.

**Figure 2:**
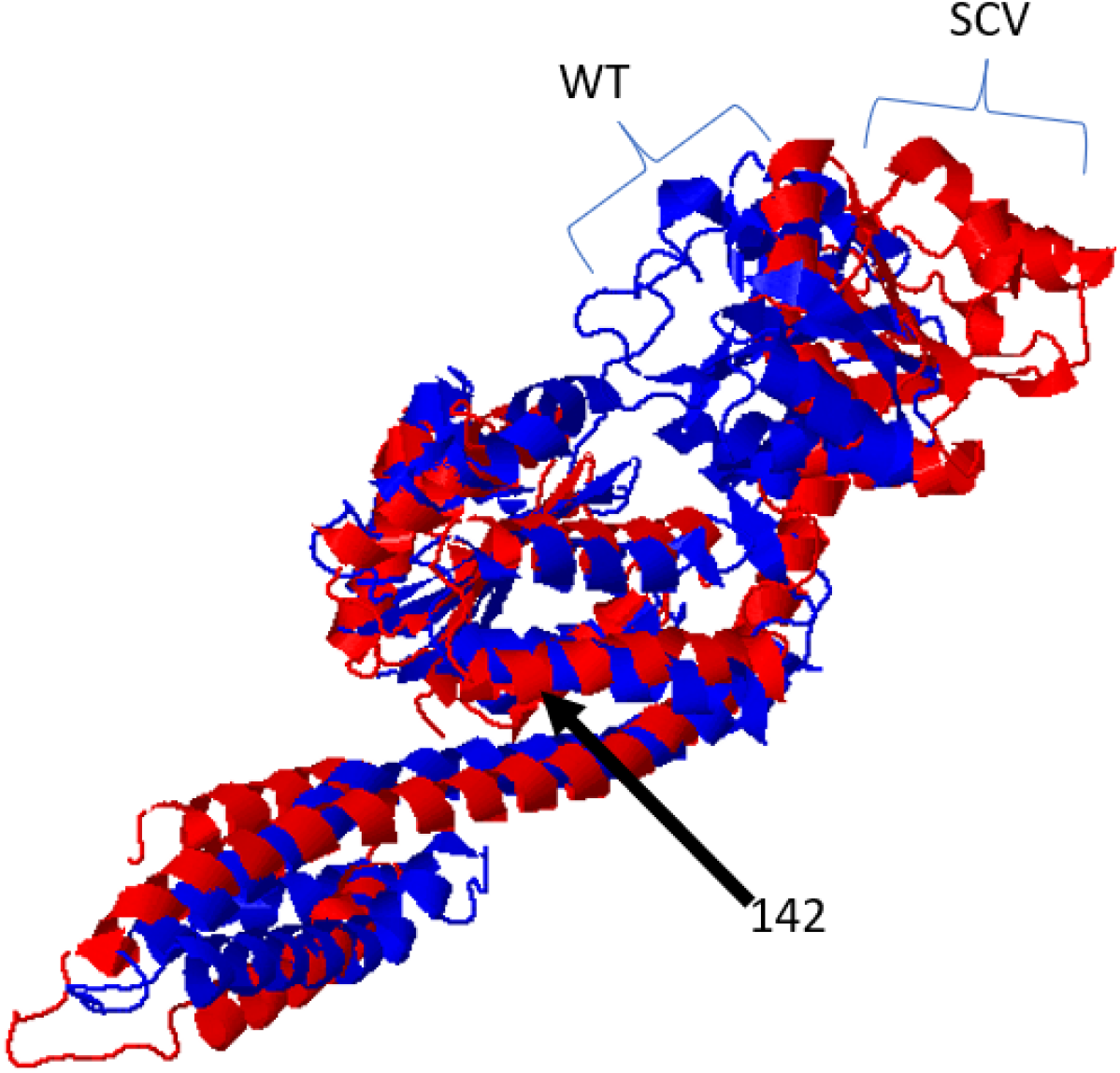
Structural alignment of WT (blue) and SCV (red) HemA. The changed amino acid at position 142 is indicated within an α-helix. The differential predicted position of the tertiary structure of the HemA is indicated with brackets.

### Confirmation the role of hemA in the small colony variant phenotype

As HemA is involved in the biosynthetic pathway of haem, we hypothesised that *E. coli* 10129 LB_2.6 would be auxotrophic for haem due to the mutation in *hemA*. Therefore, we supplied hemin exogenously during growth on agar. We found that there was a reversion of the SCV phenotype to normal size of the *E. coli* 10129 and *E. coli* 10129 LB_2 therefore this mutation is likely to be the cause of the SCV phenotype (Fig. 3). To confirm this, we rescued the SCV phenotype by transforming a plasmid with and without the cloned *hemA* from *E. coli* 10129 into *E. coli* 10129 LB_2.6. We found a reversion to a “normal” colony size in *E. coli* 10129 LB_2.6::pHSG396:hemA, while *E. coli* 10129 LB_2.6::pHSG396 did not (Fig. 4), therefore the SNP at position amino acid position 142 in *hemA* was confirmed to be responsible for a SCV phenotype in *E. coli*.

**Figure 3:**
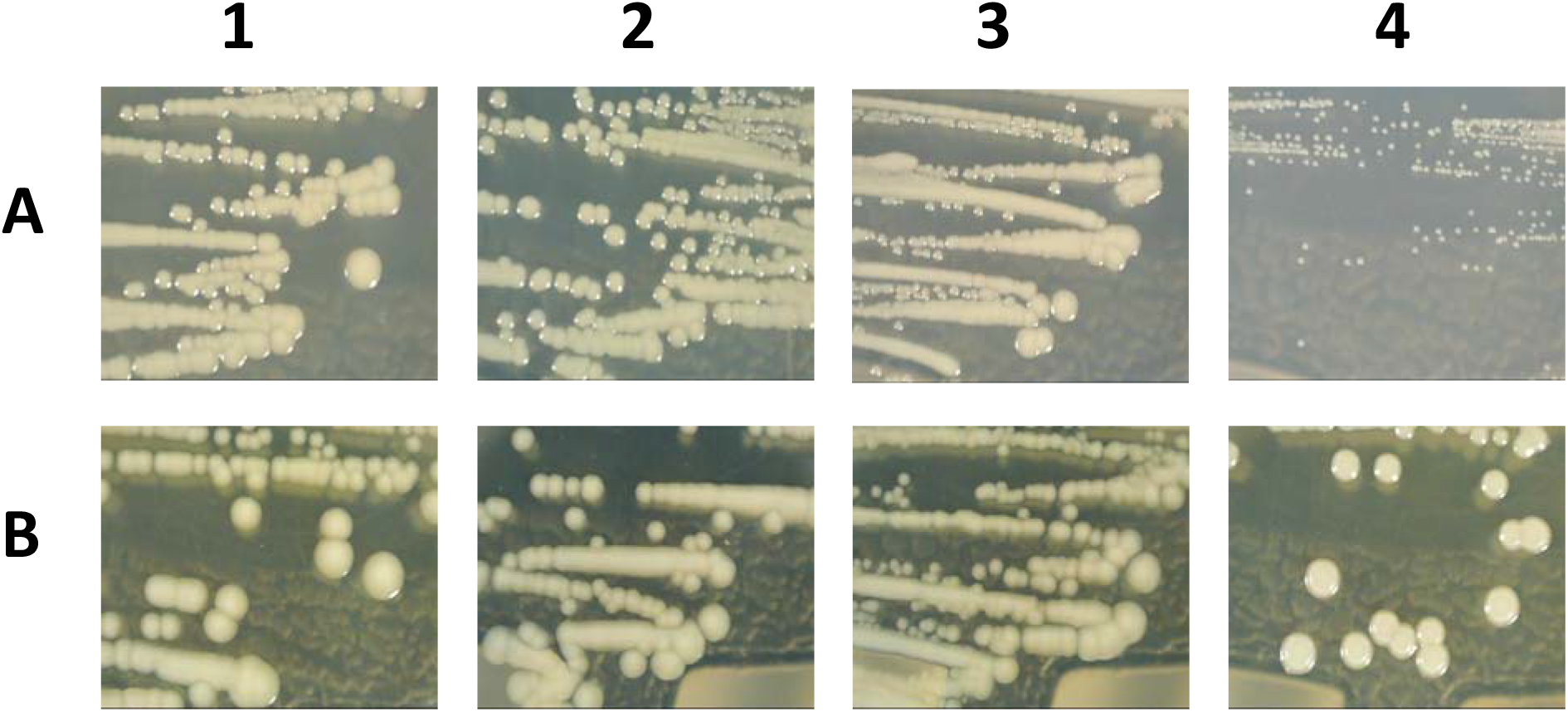
Photographs of the colonies of **1)** *E. coli* 10129, **2)** *E. coli* 10129 LB_2, **3)** *E. coli* 10129 LB_2.5 and **4)** *E. coli* 10129 LB_2.6 (SCV) grown on **A)** LB agar and **B)** LB agar supplemented with 20 μg/ml hemin

**Figure 4:**
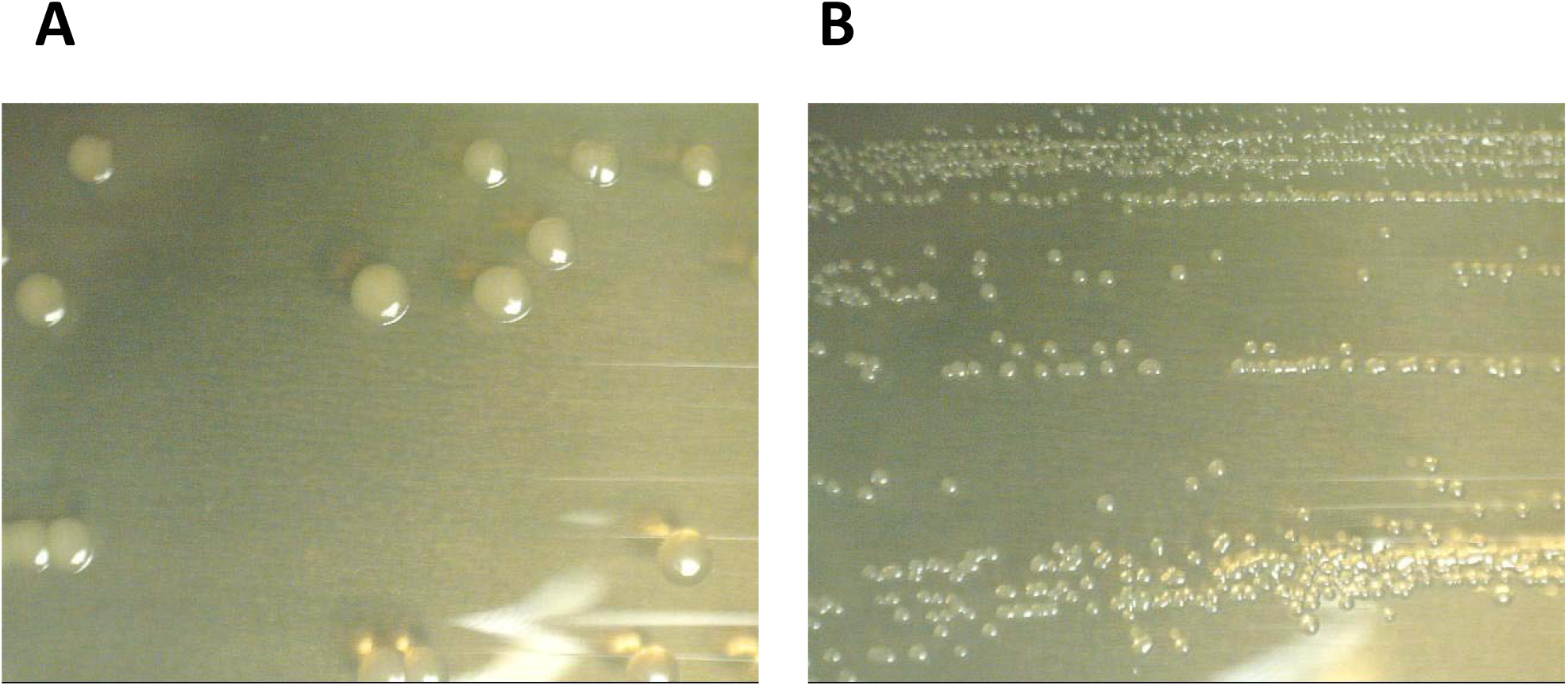
Photographs of the colonies of *E. coli* 10129 LB_2.6 transformed with **A)** pHSG396:hemA and **B)** pHSG396

### Stability of the small colony variant phenotype

*We* sought to determine whether *E. coli* 10129 LB_2.6 could revert to back to a “normal” sized colony after multiple generations. Following 40 passages on LB agar, we found no evidence of reversion of the SCV phenotype to a “normal” sized colony (Fig. 5). After 40 passages, the SNP in *hemA* was still present in *E. coli* 10129 LB_2.6 and as determined by PCR and sequencing.

**Figure 5:**
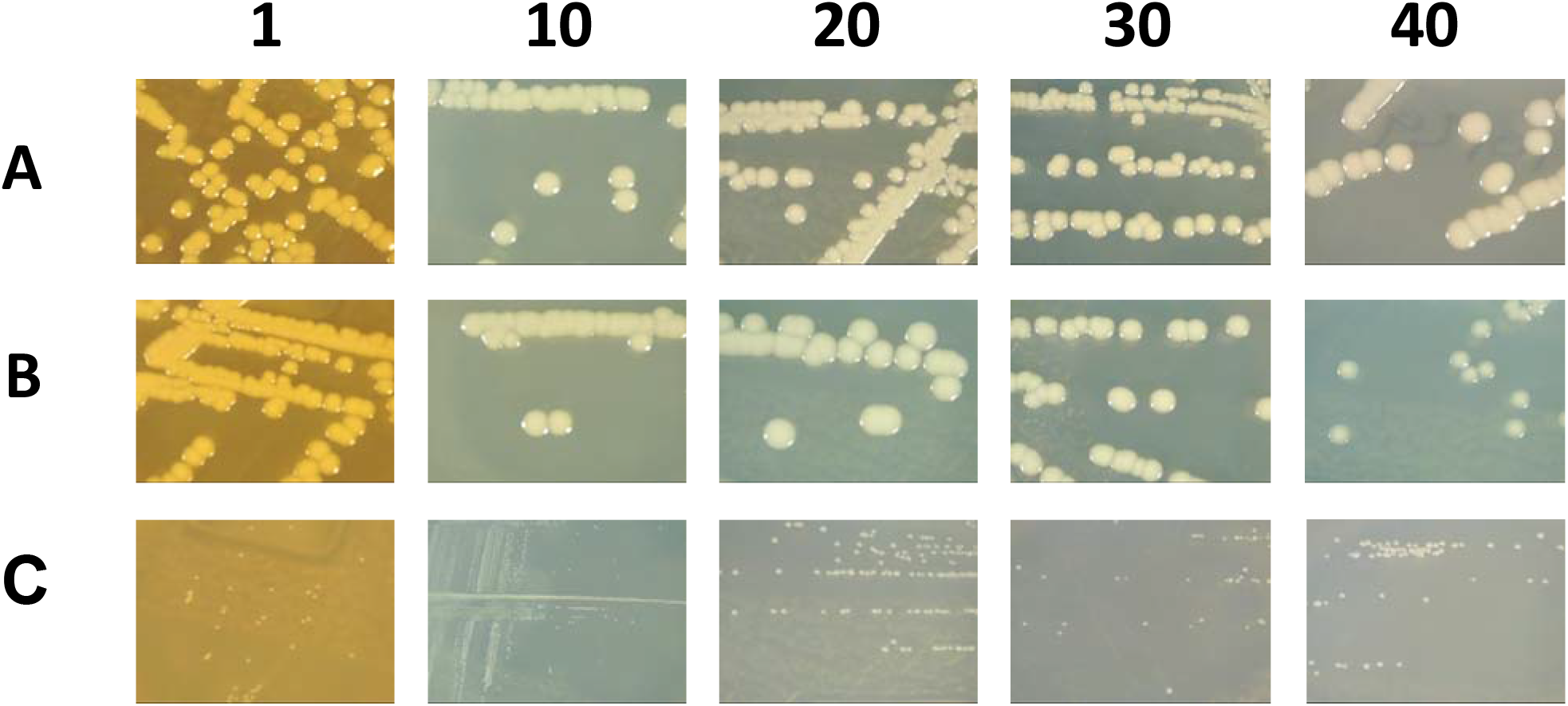
Photographs of the colonies of **A)** *E. coli* 10129, **B)** *E. coli* 10129 LB_2 and **C)** *E. coli* 10129 LB_2.6 grown on LB agar after 1, 10, 20, 30 and 40 passages

### Fitness cost of the SNP in hemA

The effect of the mutation and subsequent SCV phenotype on fitness compared to the two ancestral isolates *E. coli* 10129 and *E. coli* 10129 LB_2 was assessed competitively. We found that *E. coli* 10129 LB_2.6 carried a significant fitness cost in both LB and ISO compared to *E. coli* 10129 (LB; P value = <0.0001, ISO; P value = 0.0026) and *E. coli* 10129 LB_2 (LB; P value = <0.0001, ISO; P value = 0.0003) and, despite reversion to a “normal” sized colony on LB agar with hemin, the fitness cost imposed on *E. coli* 10129 LB_2.6 compared to *E. coli* 10129 (LB; P value = 0.0787, ISO; P value = 0.3727) and *E*.

*coli* 10129 LB_2 (LB; P value = 0.0784, ISO; P value = 0.2317) did not differ whether exogenous hemin was available (Fig. 6A). To confirm that the SNP in *hemA* was responsible for this observed fitness cost, we assessed the fitness of *E. coli* 10129 LB_2.6 and *E. coli* 10129 LB_2.6 containing either empty pHSG396 or pHSG396:hemA comparatively to the immediate ancestor *E. coli* 10129 LB_2 in LB. *E. coli* 10129 LB_2.6 containing pHSG396:hemA reverted to a relative fitness comparative to *E. coli* 10129 LB_2 (P value = 0.0877), while the relative fitness of *E. coli* 10129 LB_2.6 containing pHSG396 was found to be comparable to that of *E. coli* 10129 LB_2.6 without plasmid (P value = 0.0927) (Fig. 6B). Therefore, the SNP in *hemA* was responsible for the significant fitness cost seen in *E. coli* 10129 LB_2.6.

**Figure 6:**
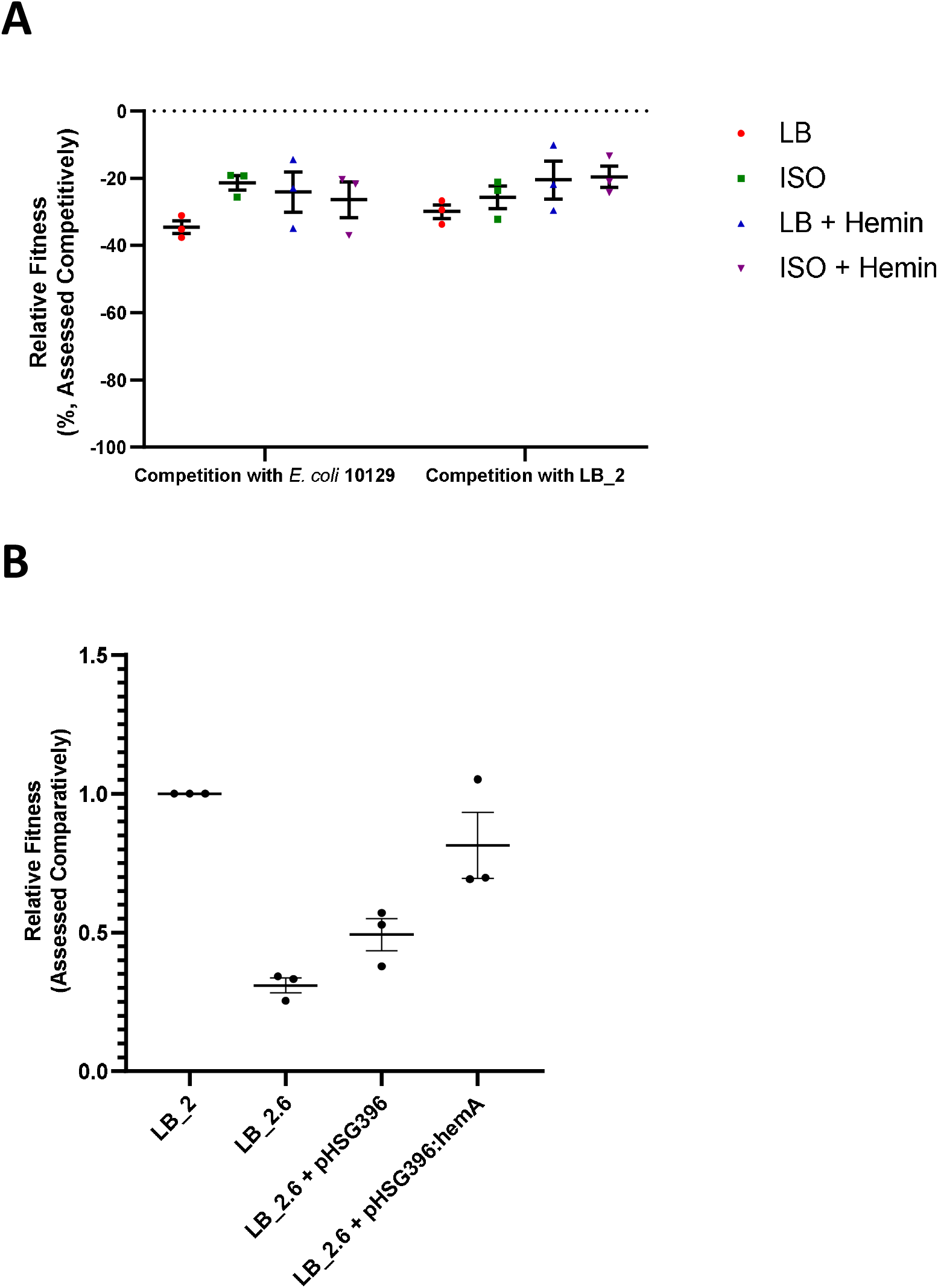
**A)** Relative fitness of *E. coli* 10129 LB_2.6 (SCV) to the immediate ancestor *E. coli* 10129 LB_2 and *E. coli* 10129 in ISO and LB, with and without addition of 20 μg/ml hemin assessed competitively. **B)** Relative fitness of *E. coli* 10129 LB_2.6, *E. coli* 10129 LB_2.6 with pHSG396 and E. coli 10129 LB_2.6 with pHSG396:hemA compared to the immediate ancestor *E. coli* 10129 LB_2, assessed comparatively.

### Biofilm production of E. coli 10129 LB_2.6

Biofilm production was significantly impaired in *E. coli* 10129 LB_2.6 compared to the immediate ancestor *E. coli* 10129 LB_2 (P value = 0.0189) when assessed in M9. However, when biofilm production was assessed in M9 supplemented with hemin none of the isolates produced a biofilm (Fig.7).

**Figure 7:**
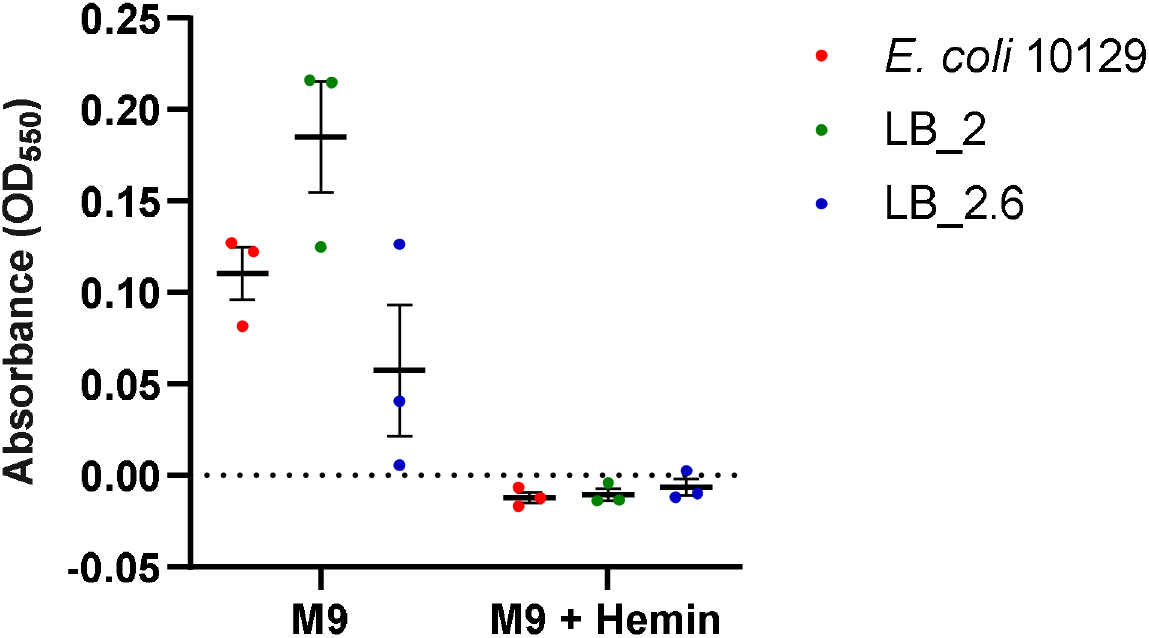
Biofilm production of *E. coli* 10129, *E. coli* 10129 LB_2 and *E. coli* 10129 LB_2.6 (SCV) in M9 and M9 supplemented with 20 μg/ml hemin.

## Discussion

Previous studies have found that SCVs of different bacterial genera have either been developed following selection in, or show resistance to, aminoglycosides [3–5] and in this study we have identified a novel SNP in *hemA* in a clinical isolate of *E. coli* which has resulted in a SCV phenotype following selection in gentamicin. Previous SNPs in *hemA* resulting in the SCV phenotype have been identified which resulted in amino acid changes at position valine-45-glycine [3,12], serine-232-stop [3,12] and glutamine-225-stop [13,14], with the latter two resulting in an early termination of the protein. The SNP we identified in this *E. coli* isolate resulted in an amino acid substitution at position 142 and did not lead to a premature stop. Mutations in *hem* genes often result in the SCV becoming auxotrophic for hemin [14, 26, 27], and the SNP we identified in *hemA* was confirmed to result in a haem auxotroph phenotype which also carries a significant fitness cost, which persists even in the presence of exogenous hemin. The SCV also has impaired biofilm formation compared to the ancestral isolates, which again did not alter in the presence of exogenous hemin. In fact, in the presence of hemin, all isolates tested in this study were unable to form biofilms.

Even though HemA is non-essential for growth, it has been identified as a potential drug target against *Acinetobacter baumannii* as it is not present in mammalian cells [28]. As the mutation in *hemA* significantly affects the fitness and biofilm production of this *E. coli* isolate, HemA may have target potential for novel compounds which could reduce fitness of *E. coli*, leading to them being outcompeted by a resident microbiome, and reduced biofilm formation which, for *E. coli*, are a common cause of catheter-associated UTIs [16].

## Conclusions

A novel SNP in *hemA* of a clinical isolate of *E. coli* results in a SCV phenotype which is auxotrophic for haem and carries a significant fitness cost even in the presence of exogenous hemin. The SCV also has impaired biofilm production relative to the ancestral isolate, opening up therapeutic possibilities with the development of *hemA* inhibitors.

## Data availability

SPAdes assembly of *E. coli* 10129 LB_2.6 was submitted to GenBank under the accession number JAAUVJ000000000.

